# Single-shot autofocus microscopy using deep learning

**DOI:** 10.1101/587485

**Authors:** Henry Pinkard, Zachary Phillips, Arman Babakhani, Daniel A. Fletcher, Laura Waller

## Abstract

Maintaining an in-focus image over long time scales is an essential and non-trivial task for a variety of microscopic imaging applications. Here, we present an autofocusing method that is inexpensive, fast, and robust. It requires only the addition of one or a few off-axis LEDs to a conventional transmitted light microscope. Defocus distance can be estimated and corrected based on a single image under this LED illumination using a neural network that is small enough to be trained on a desktop CPU in a few hours. In this work, we detail the procedure for generating data and training such a network, explore practical limits, and describe relevant design principles governing the illumination source and network architecture.

---

Many biological experiments involve imaging samples on a microscope over time periods of hours or days, either to observe the same structure over time, or to generate large tiled images by scanning a sample and stitching together a large field-of-view (FoV). In the former scenario, thermal fluctuations can induce focus drift, and in the latter, a sample that is not sufficiently flat necessitates refocusing at each position. Since it is often experimentally impractical or cumbersome to manually maintain focus, an automatic focusing mechanism is essential.

A variety of hardware and software solutions have been developed for autofocus. Broadly, these methods can be divided into two classes: hardware-based schemes that attempt to directly measure the distance from the objective lens to the sample [1, 2, 6, 4, 21], and software-based methods that take one or more out-of-focus images and use them to determine the optimal focal position [20, 9, 8]. The former usually require hardware modifications to the microscope (e.g. an infrared laser interferometry setup, additional cameras or optical systems), which can be expensive and place constraints on other aspects of the imaging system. Software-based methods, on the other hand, can be slow. For example, a software-based method might require a full focal stack, then use some measure of image sharpness to compute the ideal focal plane. More advanced methods attempt to reduce the number of images needed to compute the correct focus to just a single out-of-focus image (single-shot autofocus). However, existing single-shot methods either rely on nontrivial hardware modifications such additional lenses and sensors [8] or are limited in their application to specialized defocus regimes (i.e. can only correct defocus in one direction and in a certain range) [9]. Here, we demonstrate a new software-based single-shot autofocus that does not suffer from the limitations of previous methods. Specifically, the only hardware modification it requires is the addition of one or more off-axis light-emitting diodes (LEDs) as an illumination source, and it can correct defocus based on a single out-of-focus image.

The central idea of our method is that a neural network can be trained to predict focus from a single image taken under coherent or nearly coherent illumination at arbitrary focus relative to the sample. Data were collected using a Zeiss Axio Observer microscope (20 × 0.5 NA ob jective lens) with the illumination source replaced by a quasi-dome LED-array [13]. To train and validate our method, focal stacks were collected using Micro-Magellan [14] as software control of the microscope. Focal stacks had a total range of 60 *μ*m with 1 *μ*m spacing between successive images and were collected on the same part of the sample with two different types of illumination: one for computing the ground truth focal position from an entire stack, and the other for training the network to predict the output of this computation from a single image (Fig. 1a). The former was achieved with images taken under relatively incoherent, asymmetric illumination in order to create phase contrast from otherwise transparent cells [18]. This was achieved by using the LED array to project a half annulus source pattern. The latter was achieved by illuminating the sample with a single off-axis LED. Image sharpness was computed for each image in the incoherent focal stack by summing the high-frequency content of the image’s power spectrum, and the maximum of the resultant curve was used to determine ground truth focal position for the stack (Fig. 1a, left). We found empirically that incoherent illumination worked much better for this purpose than coherent illumination, likely due to out-of-focus blurring. Because this ground truth value is calculated by a deterministic algorithm, this paradigm scales well to large amounts of training data. Only one coherent image is needed per a training example, but we collect an entire stack to get a set of training examples equal to the number of focal planes. The defocus prediction network architecture (Fig. 1a, right) begins with a single coherent image. This image is Fourier transformed, and the magnitude of the complex-valued pixels in the central part of the Fourier transform are reshaped into a single vector. This vector is normalized to have unit mean to account for differences in illumination brightness, and it is then used as the input layer of a neural network trained in TensorFlow [3]. After the network has been trained, it can be used to correct defocus during an experiment by capturing a single image at an arbitrary defocus under the same coherent illumination, using the network to predict defocus distance, then moving to the correct focal position (Fig. 1b). A single prediction from a 2048×2048 image takes ~50 ms on a desktop CPU.

**Figure 1:**
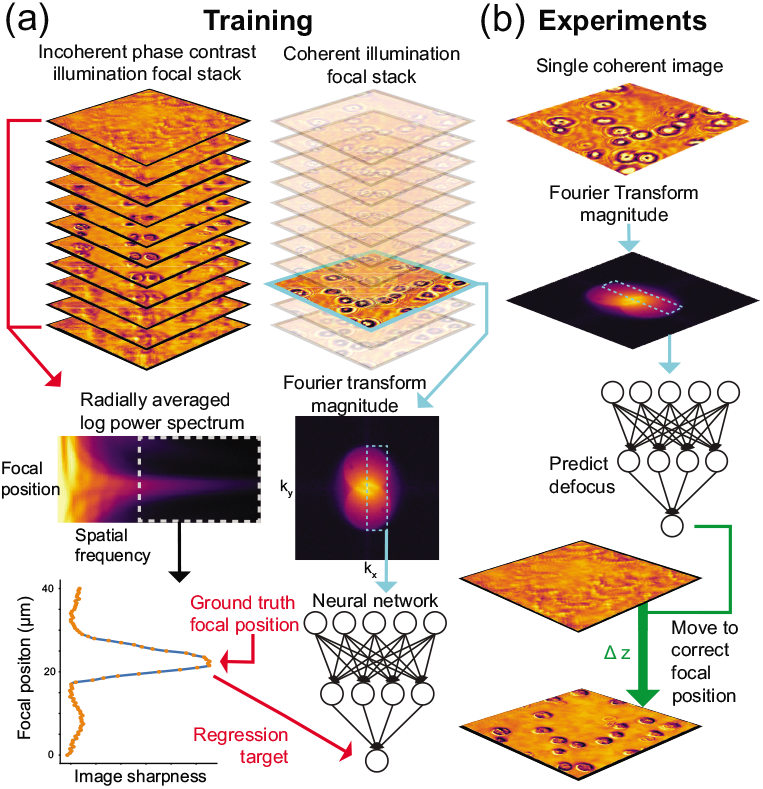
Training and defocus prediction. a) Training data consists of two focal stacks for each part of the sample, one with incoherent phase contrast, and one off-axis coherent illumination. (Left) The high spatial frequency part of each image’s power spectrum from the incoherent stack is used to compute a ground truth focal position. (Right) A single coherent image is Fourier transformed, and the magnitude of the central pixels are used as input for a neural network that is trained to predict defocus. This process is repeated for each of the coherent images in the stack to generate a set of training examples. b) During an experiment, a single coherent image is collected off-focus and fed through the same pipeline to predict defocus.

Through experimentation, we settled on a neural network architecture consisting of 10 fully-connected hidden layers of 100 units each followed by a single scalar output (i.e. the defocus prediction). In addition, we experimented with several other hyperparameters to improve training time and generalization ability. The most successful of these were: 1) applying dropout [16] to the vectorized Fourier transform input layer (but not other layers) 2) Dividing the input image into patches, and averaging the predictions over each patch 3) Using only the central part of the Fourier transform magnitude as an input vector. Each of these were manually tuned to maximize performance.

Using this architecture, we were able to train networks capable of predicting defocus with rootmean-squared error (RMSE) smaller than the axial thickness of the sample (cells). Training with 440 focal stacks took 1.5 hours on a desktop CPU or 30 minutes on a GeForce GTX 1080 Ti GPU, in addition to 2 minutes per focal stack for precomputing ground truth focal planes and Fourier transforms.

To test the performance of our method for different samples, we collected data on two different types of samples: white blood cells attached to coverglass and an unstained, mounted 5 *μ*m thick histology section (Fig. 2a). Figure 2b shows the performance of defocus predictions on a validation set based on the number of focal stacks used to train the network, where each focal stack contained 60 planes spaced 1 *μ*m apart, distributed symmetrically around the true focal plane. This curve can be quite different depending on the sample type and quality of training data. In general, we observed better performance training on noisier and more varied inputs (i.e. cells at different densities, particularly lower densities, and different exposure times). This is consistent with other results in deep learning, where adding noise to training data improves performance [19].

**Figure 2:**
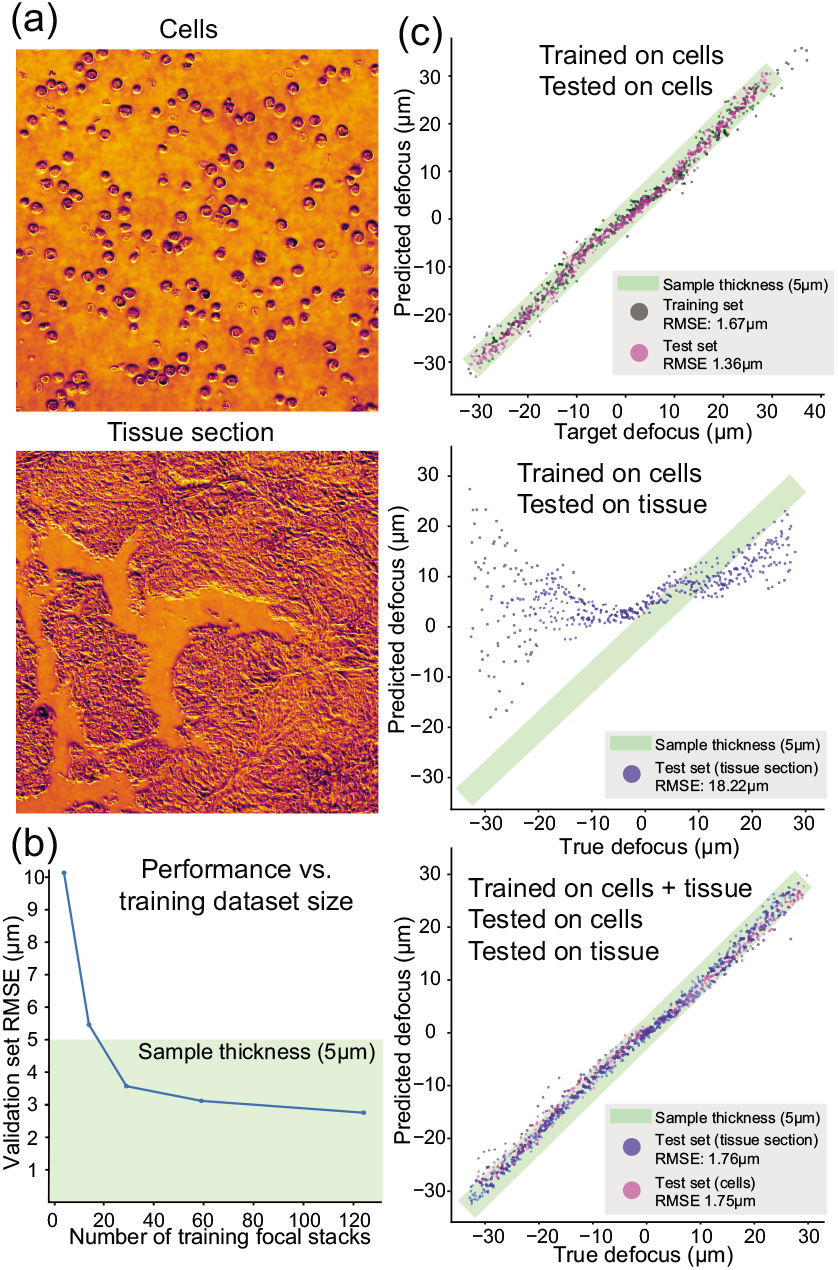
Performance across sample types. a) Representative images of cells and tissue section samples. b) Defocus prediction performance (measured by validation RMSE) as a function of the number of focal stacks used during training. c) A network trained on focal stacks of cells predicts defocus well in other samples of cells, but fails at predicting defocus in tissue sections. With additional training on limited tissue section data, however, we can learn to predict defocus in both sample types.

Figure 2c shows the ability of the network to generalize to new sample types. It performs well on different samples of the same type (i.e. trained on one slide of blood cells and tested on another slide of blood cells). Training on cells, then testing on a different type of sample (tissue section) yields poor performance. However, we can generalize to other sample types by diversifying the training data. Additional training using a smaller amount of data from the new sample type (here 130 focal stacks, compared to 440 stacks of cell data it was originally trained on) is sufficient. The best performing neural networks in other domains are typically trained on large and varied datasets [7]. If this architecture is trained on defocus data from a variety of sample types, it should generalize to new types more easily.

Empirically, we discovered that discarding the phase of the Fourier transform and using only the magnitude as the input to the network dramatically boosted performance. This observation is further supported by comparing networks trained using the Fourier transform magnitude as input vs. those trained on the argument of the Fourier transform phase (Fig. 3a). Not only were networks using magnitude able to better fit the training data, they also generalized better to a validation set. This suggests useful information for predicting defocus in a coherent intensity image is relatively more concentrated in the magnitude compared to the phase of its Fourier transform.

**Figure 3:**
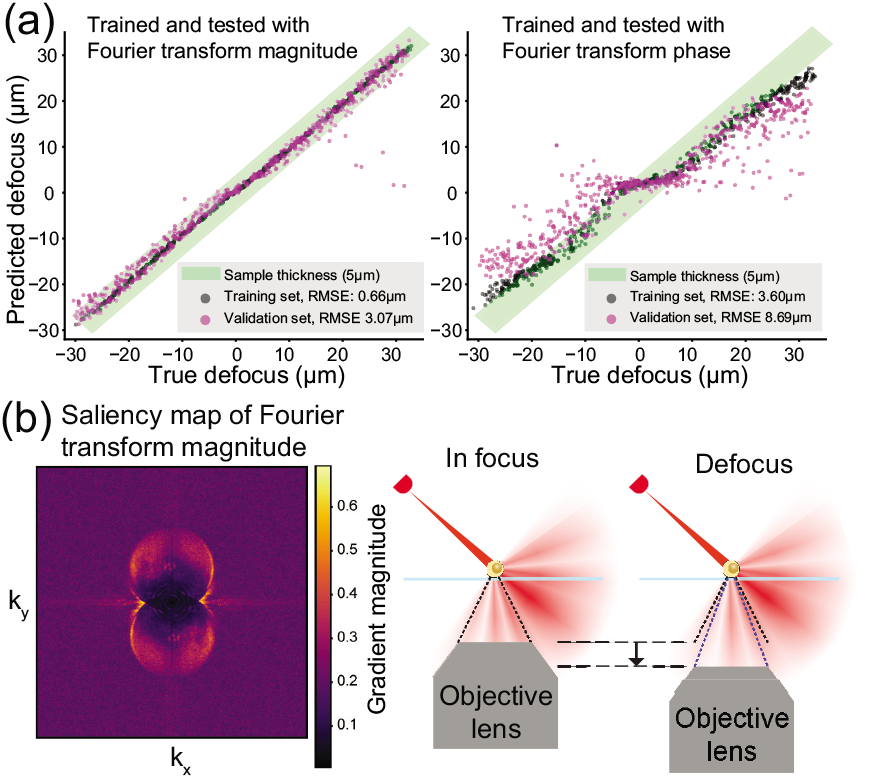
Understanding how network predicts defocus. a) A network trained on the magnitude of the Fourier transform performs better than one trained on the argument of phase of the Fourier transform. b) Left, a saliency map (the magnitude of the defocus prediction’s gradient with respect to the Fourier transform magnitude) shows the edges of the object spectrum have the strongest influence on defocus predictions. Right, edges correspond to high-angle scattered light, which may not be captured off-focus.

In order to understand what features of the images the network learns to make predictions from, we compute a saliency map. The saliency map attempts to identify which parts of the input the neural network is using to make decisions, by visualizing the gradient of a single neuron with respect to the input [15]. The idea is that the output is more sensitive to features with a large gradient and thus these have a greater influence on prediction. In our case, the gradient of the output neuron (i.e the defocus prediction) was computed with respect to the the Fourier transform magnitude. Averaging the magnitude of the gradient image over many examples clearly shows that the network recognizes specific parts of the the overlapping two-circle structure [5] that is typical for an image formed by coherent off-axis illumination (Fig. 3c). In particular, regions on the edges of the circles have an especially large gradient. These areas correspond to the highest angles of light collected by the objective lens. Intuitively, this makes sense because changing the focus will lead to proportionally greater changes in the light collected at the highest angles (Fig. 3c).

Finally, we analyze the choice of illumination on our defocus prediction network performance. Since our setup has a programmable illuminator [13], we can choose the source patterns at will. First, using one LED at a time, we tested how the angle of single-LED illumination affected performance (Fig. 4a). We found that performance improves with increasing angle of illumination, up to a point where performance rapidly degrades. This drop-off occurs in the ‘darkfield’ region (where the illumination angle is larger than the objective’s NA), likely due to the low signal-to-noise ratio (SNR) of the higher-angle darkfield images (see inset images in Fig. 4a). This drop in SNR could plausibly be caused by either a decrease in the number of photons hitting the sample from higher angle LEDs, or a drop off in the content of the sample itself at higher frequencies. To rule out the first possibility, we compensated for the expected number of photons incident on a unit area of the sample, which is expected to fall off approximately proportional to 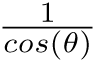, where *θ* is the angle of illumination relative to the optical axis [12]. The dataset used here increases exposure time in proportion to cos(*θ*) in order to compensate for this. Thus, the degradation of performance at high angles is most likely due to the amount of high frequency content in the sample itself at these angles and therefore might be somewhat sample-specific.

**Figure 4:**
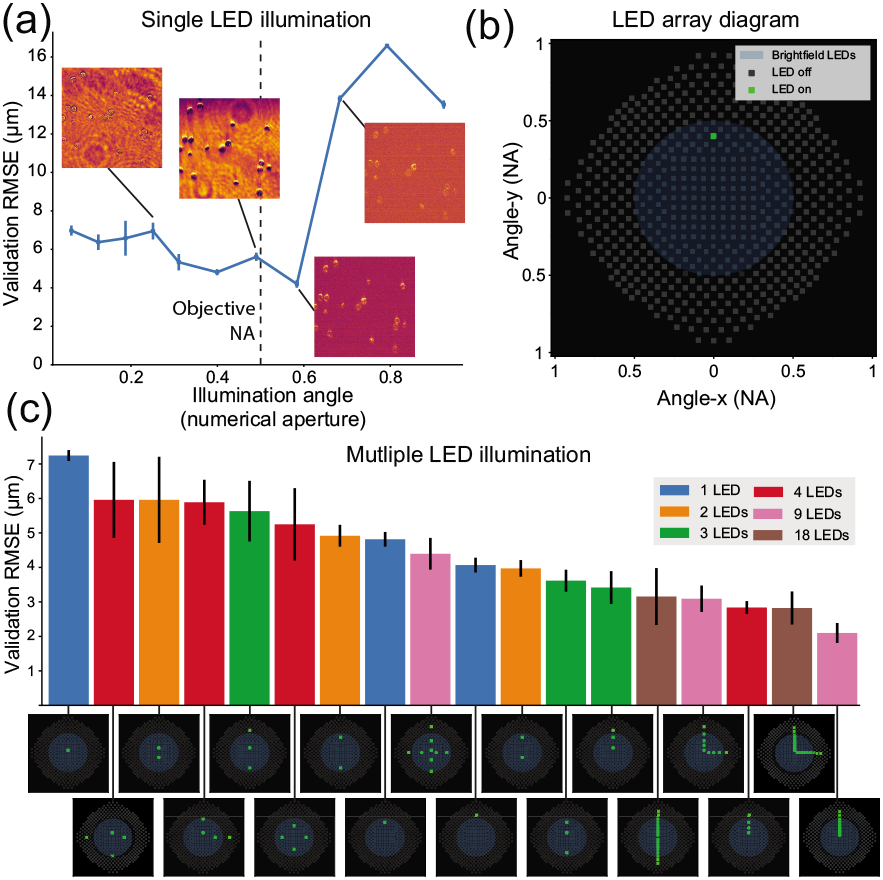
Illumination design. a) Increasing the numerical aperture (NA) (i.e. angle relative to the optical axis) of single-LED illumination increases the accuracy of defocus predictions, up to a point at which it degrades. b) Diagram of LED placements in NA space for our LED quasidome. c) Defocus prediction performance for different illumination patterns. Patterns with multiple LEDs in an asymmetric line show the lowest error.

Finally, we tested 18 different source patterns chosen from within the distribution of *x* and *y* axis-aligned LEDs available on our quasi-dome (Fig. 4b). Since the light from any two LEDs is mutually incoherent, single-LED images can be added digitally to synthesize the image that would have been produced with multiple-LED illumination. This enabled us to computationally experiment with different illumination types on the same sample. Figure 4c shows the defocus prediction performance of various patterns of illumination. The best performing patterns were those that contained multiple LEDs arranged in a line. Given that specific parts of the Fourier transform contain important information for defocus prediction and that these areas will move to different parts of Fourier space with different angles of illumination, we speculate that the line of LEDs helps to spread relevant information for defocus prediction into different parts of the spectrum. Although this analysis demonstrates more and higher angle LED patterns seem to yield superior performance, there are potential caveats: In the former case, it could fail to hold when applied to a denser sample (i.e. not a sparse distribution of cells). In the latter, there is the cost of the increase in exposure time needed to acquire such images.

To summarize, we have demonstrated a method for training and using a neural network for single-shot autofocus, with analysis of design principles and practical trade-offs. The method works with different sample types and is simple to implement on a conventional transmitted light microscope, requiring only the addition of off-axis illumination. Alternately, it can be thought of as a new modality for existing coded-illumination setups, which have been demonstrated for super-resolution [22, 11, 17], quantitative phase [22, 18, 11], and multi-contrast microscopy [23, 10]. Our method provides a low-cost, simple mechanism for automated autofocus if coded illumination becomes standardized on microscopes of the future. Additionally, we have shown an example of how deep learning can be mixed with conventional signal processing techniques to boost performance and and interpret the functionality of neural networks.

## Open source

The code needed to implement this technique and reproduce all figures in this manuscript can be found in the Jupyter notebook: 1. H. Pinkard, “Single-shot autofocus microscopy using deep learning–code,” (2019), https://doi.org/10.6084/m9.figshare.7453436.v1. Due to its large size, the corresponding data is available upon request.

## Funding and acknowledgements

This project was funded by Packard Fellowship and Chan Zuckerberg Biohub Investigator Awards to Laura Waller and Daniel Fletcher, STROBE: A National Science Foundation Science & Technology Center under Grant No. DMR 1548924 awarded to Laura Waller, a National Institutes of Health R01 grant to Daniel Fletcher, a National Science Foundation Graduate Research Fellowship awarded to Henry Pinkard, and a Berkeley Institute for Data Science/UCSF Bakar Computational Health Sciences Institute Fellowship awarded to Henry Pinkard with support from the Koret Foundation, the Gordon and Betty Moore Foundation through Grant GBMF3834 and the Alfred P. Sloan Foundation through Grant 2013-10-27 to the University of California, Berkeley.

The authors thank S.Y. Liu for providing the tissue sample and BIDS and its personnel in providing physical space, general logistical and technical support.

